# Detection of a hybrid PrPfr state in the dark reversion of a bathy phytochrome indicates inter-dimer allostery

**DOI:** 10.1101/2025.05.28.656462

**Authors:** Sayan Prodhan, Petra Mészáros, Szabolcs Bódizs, Yalin Zhou, Micha Maj, Sebastian Westenhoff

## Abstract

Phytochromes are photosensor proteins which detect light in plants, fungi, and bacteria. They photoswitch between red light absorbing (Pr) and far-red light absorbing (Pfr) states, however, thermal reversion in the dark is an equally important factor in controlling their signaling levels. Phytochromes are generally dimeric proteins, and mixed PrPfr states are therefore possible. These states have been implied in the dark reversion studies of plant phytochromes, but not in bacterial phytochromes. Here, we investigate the dark reversion kinetics of the ‘bathy’ phytochrome from *P. aeruginosa* (*Pa*BphP) using UV-Vis absorption spectroscopy. A single set of time-resolved spectra does not conclusively reveal the presence of a mixed PrPfr state, as both a direct Pr *→* Pfr model or a sequential Pr *→* PrPfr *→* Pfr model fit the spectral kinetics. However, a systematic analysis of dark reversion kinetics with varying Pr/Pfr ratios can only be satisfactorily fit by the sequential model, which indicates the presence of an intermediate PrPfr state. A newly designed monomeric variant of *Pa*BphP provides strong support for this interpretation. Temperature-dependent kinetics revealed similarly low activation energies for the dark reversion processes of both proteins, consistent with a previously proposed keto-enol tautomerization preceding dark reversion. Interestingly, our results suggest allosteric regulation of dark reversion across the dimer, which we propose to be a contributing factor in phytochrome signaling.

## INTRODUCTION

Many organisms rely on photoreception to adapt to changing environmental conditions ^1,2^. Among various light-sensing systems, phytochromes are red light-detecting photoreceptors that regulate physiological responses in plants, fungi, and bacteria ^3–5^. In plants, they control essential processes like seed germination, chloroplast development, and flowering ^6–8^, while in bacteria, they regulate pigmentation, motility, and biofilm formation ^9–11^. Therefore, the phytochrome photocycle is extensively studied for biotechnological applications, as well as for the engineering of optogenetic tools and near-infrared fluorescent markers used in microscopy and cell biology ^12,13^.

Phytochromes are typically homodimeric proteins with a modular structure ^14^. Each monomer consists of an N-terminal photosensory core module (PCM), responsible for light sensing, and a C-terminal output module (OPM), which mediates signal transduction. In some cases, additional N-terminal extensions may also contribute to the effector function of the protein. The PCM is composed of three functionally distinct domains: PAS (Per-ARNT-Sim) ^15^, GAF (cGMP phosphodiesterase / adenylate cyclase / FhlA) ^16^, and PHY (phytochrome-specific domain), and contains a bilin cofactor that serves as the chromophore ^17^.

Phytochromes interconvert between two structurally distinct states: the red light-absorbing Pr state (*λ*_max_ *≈* 700 nm, for bacterial phytochromes) and the far-red-light-absorbing Pfr state (*λ*_max_ *≈* 750 nm). These states are associated with distinct biochemical properties and, consequently, different signaling activities ^18,19^. Based on the relative free energies of these states, phytochromes are classified as either canonical, the most common type with a thermally stable Pr state, or reverse-activated (bathy), in which the Pfr state is thermodynamically favored (Fig.1(a)) ^20–22^. In the absence of light, phytochromes thermally revert to their resting state.

**Figure 1:**
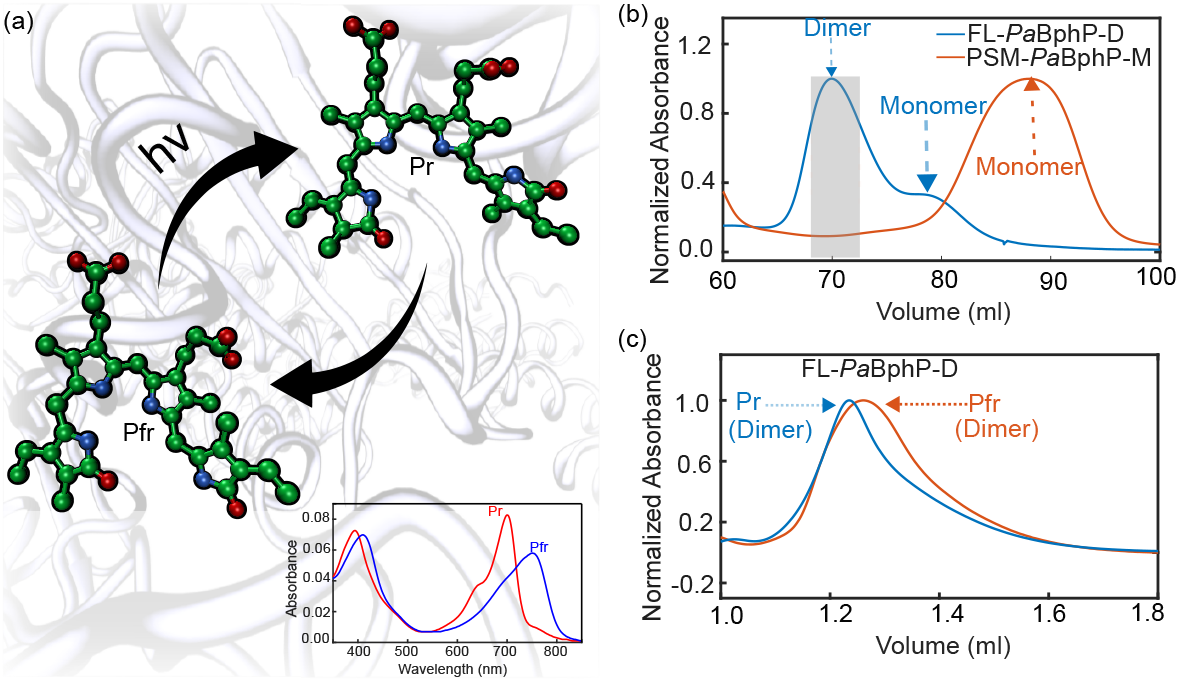
The phytochrome photoswitches between Pr and Pfr states, and chromatography confirms the oligomeric states of the two protein samples. (a) Conformational differences in the biliverdin structure distinguish the Pfr (≈ 750*nm*) and Pr (≈ 700*nm*) states, which are reflected in their steady-state absorption spectra.(b) Preparatory SEC analysis of FL-*Pa*BphP-D and PSM-*Pa*BphP-M revealing their oligomerization state. The shaded region indicates the fraction collected fractions for FL-*Pa*BphP-D. For PSM-*Pa*BphP-M, the collection zone was around the peak maximum. (c) Analytical SEC of FL-*Pa*BphP-D in Pr and Pfr state confirms that the protein is predominatly dimeric. The slight shift in the retention time between the states likely reflects conformational changes.

Canonical and bathy phytochromes have been extensively studied using spectroscopy and structural biology methods. Most of these studies focus on understanding the sequence of events during photoactivation ^23–28^. They show that upon photoactivation, the chromophore undergoes isomerization around the C15=C16 bond, followed by a series of structural changes through intermediate states (i.e., Lumi-F/R and Meta-F/R). ^29–31^. Crystal, NMR, and cryo-EM structures of phytochromes in both the Pr and Pfr states have been solved ^32–37^. Of particular interest are two recent crystal and cryo-EM structures, which reveal phytochromes in mixed Pr/Pfr heterodimers ^38,39^. However, it remains unclear how these intermediate states are formed. The dark reversion has been studied in both plant and bacterial phytochromes ^19,25,28,29,35,37,40–5^ with studies on plant phytochromes providing the most detailed insights, including differences in activation energy across various temperature ranges ^40^, the presence of PrPfr hybrid states ^54^, and temperaturedependent kinetics that allow phytochromes to function as temperature sensors ^19,50^. Plants also have multiple isoforms of phytochromes, which contribute to varying dark reversion rates, allowing for more complex adaptation to different environmental conditions ^43^.

In bacterial phytochromes, the dark reversion process is less well understood. While dark reversion is commonly used as a standard assay in structural and biochemical studies of phytochromes, only a few detailed studies have been conducted ^47,51,52,57–59^. However, a hybrid PrPfr state has not been observed as part of the dark reversion process in bacterial phytochromes yet.

In this study, we systematically investigate the dark reversion kinetics of a bathy phytochrome from *Pseudomonas aeruginosa* (*Pa*BphP). By comparing the wild-type full-length dimer (FL-*Pa*BphP-D) with a newly designed monomeric variant (PSM-*Pa*BphP-M), we provide clear evidence for the presence of PrPfr states as intermediates in the dark reversion process for the wild-type protein. This finding suggests a novel allosteric mechanism between the two protomers.

## MATERIALS AND METHODS

### Recombinant protein production

The plasmid used for the recombinant expression of the full-length *P. aeruginosa* phytochrome contained the *Pa*BphP insert (GenBank: AAG07504.1) with a C-terminal *His*_6_-tag and a pET28a(+) backbone (Novagen). The plasmid used for the recombinant expression of the *P. aeruginosa* phytochrome PAS-GAF-PHY mutant fragment contained the *Pa*BphP insert (GenBank: AAG07504.1) with a C-terminal His6-tag and a pET24a(+) plasmid backbone (GenScript). To facilitate biliverdin incorporation during expression, all the phytochrome constructs were co-expressed with a heme oxygenase plasmid, kindly provided by Prof. Janne A. Ihalainen. The plasmids were transformed into BL21(DE3) *Escherichia coli* strain for expression.

Cells were grown in LB medium supplemented with 50 *µ*g/mL kanamycin and 34.5 *µ*g/mL chloramphenicol at 37°C until reaching an *OD*_6_*oo* of 0.8. Expression of *Pa*BphP and heme oxygenase was induced by the addition of 1 mM IPTG and 1 mM 5-aminolevulinic acid. Protein expression was carried out overnight at 18°C.

### Protein purification

Cells were re-suspended in 10 mL of lysis buffer per gram of pellets (50 mM TrisHCl, pH 8.0, 150 mM NaCl, 5% glycerol), supplemented with one tablet of EDTA-free protease inhibitor (cOmplete Protease Inhibitor Cocktail, Roche) and 10 U/mL DNase I. Lysis was carried out by sonication on ice at 70% amplitude for 5 min, with 5 s pulses on and 10 s pulses off. The supernatant was collected after centrifugation at 20,000g for 35 min (JA 14.50 rotor, Beckman) and was briefly incubated with a molar excess of biliverdin hydrochloride.ă The lysate was purified by Ni-NTA affinity chromatography using a HisTrap HP 5 mL column (Cytiva). The bound protein was washed with wash buffer I (50 mM Tris, 50 mM NaCl, pH 8.0) and wash buffer II (50 mM Tris, 1 M NaCl, pH 8.0), and eluted with elution buffer (50 mM Tris, 50 mM NaCl, 300 mM imidazole, pH 8.0). This was followed by size-exclusion chromatography using a HiLoad 16/600 Superdex 200 pg column (Cytiva) in wash buffer I. The purified protein was then flash-frozen for storage.

### Design of monomeric variant

The mutant plasmid was designed using only the photosensory module (PSM) of the protein, excluding the histidine kinase module, to promote monomerization with minimal mutational changes. Mutation sites were selected based on amino acid properties and spatial proximity (within 4 Å). I294E and V298E mutations were introduced into the wild-type sequence to replace hydrophobic residues with charged glutamate residues (Fig. 2(a)). The mutant plasmid was generated using GenScripts site-directed mutagenesis service.

**Figure 2:**
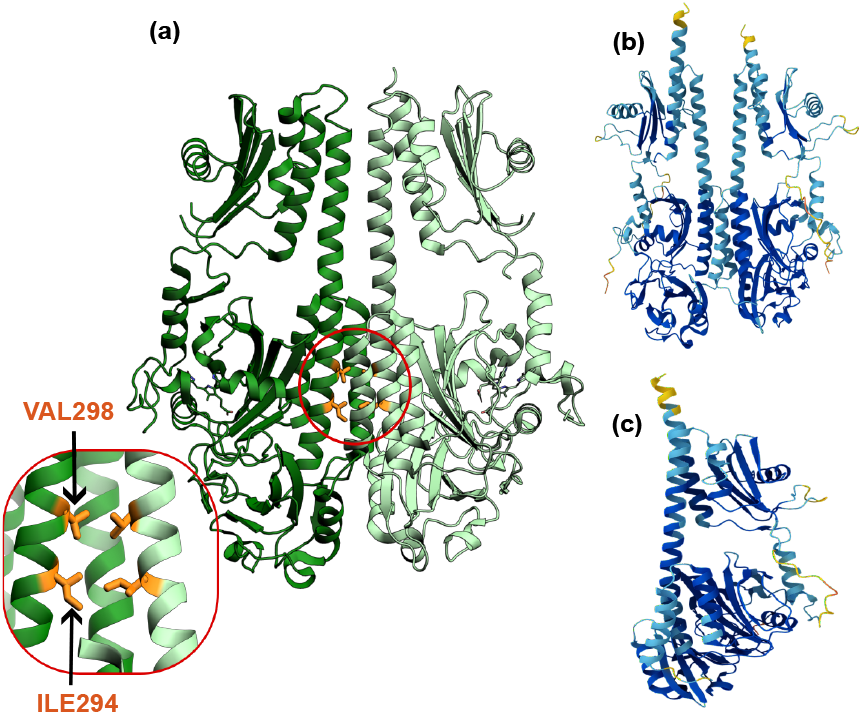
Design and structural prediction of truncated monomeric mutant *Pa*BphP construct. (a) Locations of I294E and V298E mutation sites in the wild-type, truncated structure, highlighting their spatial proximity. (b) AlphaFold prediction of the wild-type truncated sequence as a dimer, showing proper dimerization and a conformation comparable to the crystal structure. (c) AlphaFold prediction of the I294E/V298E mutant in its monomeric form, showing proper folding of the monomeric unit.

### UV-Vis absorption spectroscopy

Steady-state UVVis absorption measurements were performed using a Shimadzu UV-1900i spectrophotometer equipped with LabSolutions UVVis software (Version 1.13). For dark reversion kinetics, 100 sequential UVVis spectra scans were recorded at 45-second intervals with a delay of 4 seconds between diode illumination of the phytochrome samples and the start of each measurement. For time-dependent diode illumination experiments, samples were exposed to a 780 nm light-emitting diode (LED) for durations ranging from 6 to 60 seconds. Temperature-dependent measurements were conducted by irradiating the phytochrome samples with a 780 nm LED for 120 seconds at temperatures ranging from 10 °C to 35 °C. To obtain a fully uniform Pr state, the sample was illuminated with a 780 nm LED for 120 seconds. To obtain a fully uniform Pfr state, the sample was kept in darkness for 3 hours.

### Calculation of Pr/Pfr fraction in diode illuminated phytochrome

To determine the relative amounts of Pr and Pfr in the illuminated samples, we first acquired absorption spectra for the pure Pr and pure Pfr states. The pure Pfr spectrum was recorded after incubating the sample in the dark for approximately three hours. The pure Pr spectrum was recorded after illuminating the sample with a 780 nm LED for 2 minutes. The concentrations of Pr and Pfr in the illuminated sample were determined by fitting the initial spectrum obtained after illumination to the following equation:

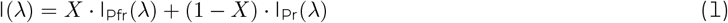

Here, I,*I*_*Pfr*_(*λ*), and *I*_*Pr*_(*λ*) are the intensity of experimental, pure Pfr and pure Pr spectra respectively. X and (1-X) represent the fractions of Pfr and Pr, respectively. We assume that no sample degradation happens during the dark reversion process.

## RESULTS

### Design of monomeric variant

To study the dark reversion kinetics as a function of the oligomeric state of PaBphP, we prepared a truncated, monomeric variant of the protein. Mutations I294E and V298E were introduced into the photosensory module (PSM) to disrupt dimer formation by weakening hydrophobic interactions and introducing electrostatic repulsion (Fig. 2(a)). This mutational strategy was motivated by previous studies on the canonical phytochrome from *Deinococcus radiodurans*, which showed that glutamate substitutions at F145, L311, and L314 positions in *Dr*BphP effectively disrupted dimerization and produced stable monomers. ^53,60^ Our AlphaFold predictions supported that these mutations would disrupt the dimer interface while maintaining proper monomer folding. Structural models of both the wild-type and mutant truncated proteins were generated using the GenBank sequence AAG07504.1, in both dimeric and monomeric forms. While the wild-type sequence was reliably predicted to form a stable dimer (Fig. 2(b)), AlphaFold did not predict a stable dimeric structure for the mutant variant. However, it successfully predicted a stable monomeric fold for the mutant (Fig. 2(c)), indicating that the introduced mutations disrupted dimerization without compromising the native folding.

### Size-exclusion chromatography

To confirm the oligomeric states of the full-length protein (FL-*Pa*BphP-D) and the truncated mutant (PSM-*Pa*BphP-M), we performed size-exclusion chromatography (SEC). Preparative SEC (Fig. 1(b)) showed that FL-*Pa*BphP-D (blue trace) predominantly forms dimers, indicated by a major peak at lower elution volumes (68-72 mL). A minor peak at 78 mL suggests a small monomeric population. The shaded region marks the fraction collected for further analysis. In contrast, PSM-*Pa*BphP-M (red trace) showed a single peak around 88 mL, consistent with monomeric species. Analytical SEC of the FL-*Pa*BphP-D fraction, performed both in the dark and under continuous 780 nm illumination, confirmed the dimeric state in both Pr and Pfr forms, with elution volumes of 1.24-1.26 mL. A small shoulder at 1.5 mL may correspond to a minor monomeric fraction (Fig. 1(c)). A slight shift in the dimer peak between states likely reflects conformational differences between Pr and Pfr that influence the proteins shape. We conclude that PSM-*Pa*BphP-M is monomeric, whereas the wild-type FL-*Pa*BphP-D is predominantly dimeric (>90%).

### Dark reversion

To measure dark reversion kinetics, all phytochrome samples were illuminated for 120 s with LED light at 780 nm to populate the Pr state. Fig. 3(a,b) shows the time-dependent spectral evolution from the initially photoconverted Pr state back to the Pfr state. Under these conditions, FL-*Pa*BphP-D and PSM-*Pa*BphP-M display qualitatively similar spectral features over time.

**Figure 3:**
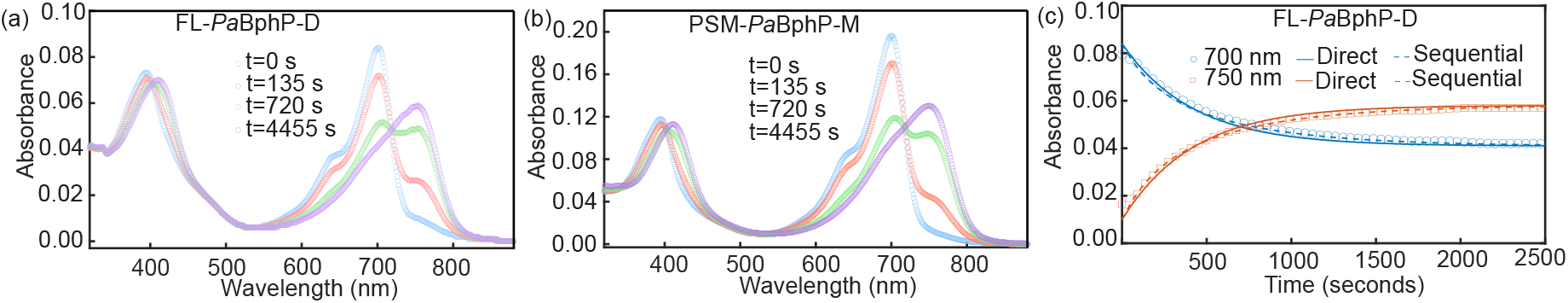
The sequential and direct model cannot be distinguished by fitting the dark reversion kinetics of FL-*Pa*BphP-D and PSM-*Pa*BphP-M. UV-Vis spectra recorded over time during dark reversion for (a) FL-*Pa*BphP-D, and (b) PSM-*Pa*BphP-M. (c) Global fitting of dark reversion kinetics for FL-*Pa*BphP-D at 700 nm and 750 nm using a direct sequential model. Solid lines represent the global fit with a direct model, while dashed lines show the fit using a sequential model. The temperature for this experiment is maintained at 30 °C.

The dark reversion kinetics of the protein is analyzed using two kinetic models to elucidate the underlying mechanism: a direct transition and a sequential model involving an intermediate state. We first describe the direct transition model:

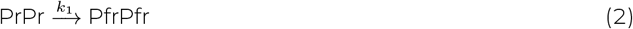

Here, *k*_1_ is the transition rate constant from PrPr to PfrPfr. The analytic solution for fitting the kinetic data is

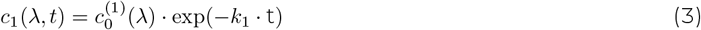

where 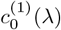 and *c*_1_(*λ, t*) are the initial and time-dependent concentrations of the PrPr state, respectively. The sequential model incorporates the presence of an intermediate state (PrPfr) between the initial and final species of the dark reversion process. This model is more plausible than the direct model for the dimeric system, provided that The model is described by:

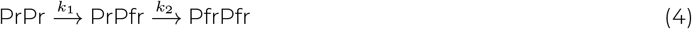

Here *k*_1_ and *k*_2_ are the dark reversion rate constant from PrPr to PrPfr, and from PrPfr to PfrPfr respectively. The solutions of the sequential model used for the global fitting are

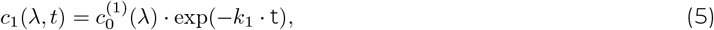

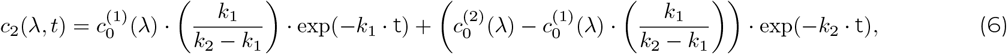

and

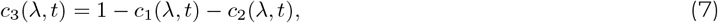

where 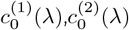 are the initial concentration of PrPr and PrPfr species. *c*_1_(*λ, t*),*c*_2_(*λ, t*), and *c*_3_(*λ, t*) represent the time-dependent concentration of the PrPr,PrPfr and PfrPfr forms, respectively. We perform global fitting of the time-dependent dark reversion spectra using these kinetic models.

In the first experiment, dark reversion kinetics is measured starting from a state with a maximal Pr population. Global fitting of the kinetic data for FL-*Pa*BphP-D is performed using both the direct (Equation 3) and sequential (Equations 5-7) models. The resulting kinetic fits at 700 and 750 nm are shown in Fig. 3(c). Although the sequential model yields a slightly better fit than the direct model, this difference is not sufficient to clearly favor one model over the other.

To differentiate between the models, we investigated the dependence of the refined rate constants on the initial photostationary state. To achieve varying initial photostationary states, we altered the duration of the illumination at 780 nm from 6 to 60 seconds. This variation in illumination time modified the initial fraction of the Pr state at the onset of the dark reversion measurement (Fig. 4(a,b)). The resulting photostationary states are defined as the fraction of Pr relative to the total phytochrome population, as given by ^44^:

**Figure 4:**
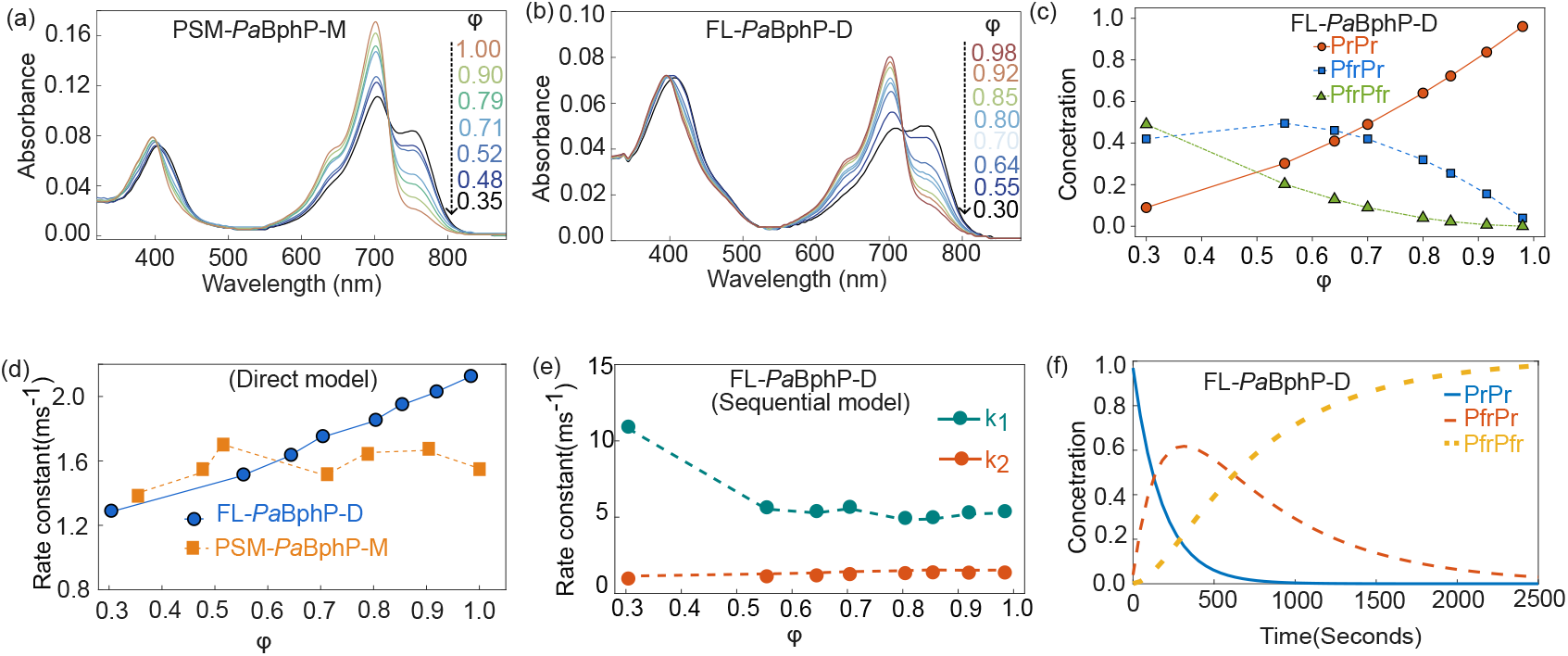
Preparation of different initial concentrations of Pr and Pfr and global kinetic fits. The temperature for this experiment is maintained at 30 °C. Spectra support the sequential model for the wild-type protein and reveal the existence of the hybrid PrPfr state. Dark reversion spectra at various photostationary states for (a) PSM-*Pa*BphP-M and (b) FL-*Pa*BphP-D. (c) Variation in the starting fractions of PrPr, PrPfr, and PfrPfr states with varying Pr concentration for the FL-*Pa*BphP-D. (d) Variation of the dark reversion rate constants, obtained from global fitting with the direct step model for FL-*Pa*BphP-D and PSM-*Pa*BphP-M. (e) Variation of the fast (*k*_1_) and slow (*k*_2_) dark reversion rate constants with Pr fraction for FL-*Pa*BphP-D. (f) Variation of simulated relative concentration of PrPr,PrPfr, and PfrPfr over time during dark reversion.

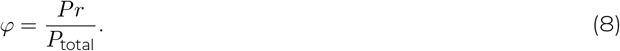

*φ* was determined by fitting the spectra with the pure spectra for Pr and Pfr (section 2.5). In terms of the sequential model, the initial relative concentrations of PrPr, PrPfr, and PfrPfr species depend on *φ* and can be calculated assuming a stochastic occupancy of the hybrid state as ^41^

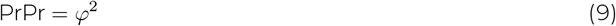

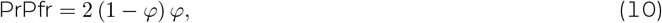

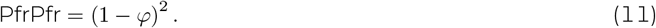

The relative concentrations of PrPr, PrPfr, and PfrPfr, needed for the global fit, are calculated using Equation 9–11 and shown in Fig. 4(c).

After global fitting of all data sets, we now analyze the rate-refined constants as a function of the initial photostationary states. Using the direct transition model (Equation 2-3) the rate constants (*k*_1_) remain independent of the initial photostationary states for PSM-*Pa*BphP-M, as shown in Fig. 4(d). This is expected as the rate constant of dark reversion should not change with the population of the initial state. In contrast, for the FL-*Pa*BphP-D, the refined rates vary significantly with *φ* (Fig. 4(d)). This dependence indicates that the direct transition model does not adequately describe the reaction, as true rate constants should be independent of initial concentrations. When examining the rates of the sequential model, refined for spectra starting from different photostationary states, we find that *k*_1_ and *k*_2_ remain constant for *φ >* 0.5 (Fig. 4(e) and Table 1). Notably, *k*_1_ at the lowest excitation fraction (*φ* = 0.3) appears as an outlier, which is likely due to the low abundance of PrPr under this illumination condition, which limits the reliable determination of *k*_1_. This analysis strongly supports the applicability of the sequential model to the dark reversion of the dimeric protein, as the rate constants are independent of the initial population distribution. The monomeric variant further confirms this, as the direct model adequately describes the dark reversion across all *φ* values.

**Table 1:**
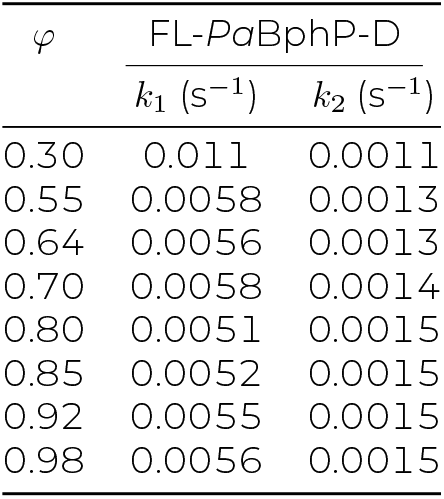
Photostationary-state-dependent dark reversion kinetic rates for FL-*Pa*BphP-D construct.

**Table 2:** Temperature dependent rate constants for FL-*Pa*BphP-D and PSM-*Pa*BphP-M construct.

| Temperature<br>(°C) | FL-*PaBphP*-D |  | PSM-*PaBphP*-M |
| --- | --- | --- | --- |
| | $k_1$ (s<sup>-1</sup>) | $k_2$ (s<sup>-1</sup>) | $k_1$ (s<sup>-1</sup>) |
| 10 | 0.0062 | 0.00168 | 0.0009 |
| 15 | 0.0064 | 0.00175 | 0.0011 |
| 20 | 0.0066 | 0.00179 | 0.0013 |
| 25 | 0.0065 | 0.00191 | 0.0014 |
| 30 | 0.0069 | 0.00191 | 0.0015 |
| 35 | 0.0067 | 0.0021 | 0.0016 |

We extended our investigation by probing the temperature dependence of dark reversion kinetics. Spectra for both proteins were recorded across a temperature range of 10°C to 35°C. Each measurement was initiated from pure Pr (PrPr) state at maximum photoexcitation. The initial spectra recorded immediately after illumination are consistent across all temperatures, confirming a uniform starting population of the Pr state. To extract the temperature-dependent rate constants, we globally fit the kinetic data using the sequential model (Equation 4) for FL-*Pa*BphP-D and the direct model (Equation 2) for PSM-*Pa*BphP-M.

The resulting temperature dependence of the rate constants is shown in Fig.5a,b and tabulated in Table 2. For both proteins, the kinetic rate constants increase significantly with temperature, indicating that thermal energy facilitates dark reversion. To quantify this effect, the activation energy (*E*_*a*_) for each kinetic step was determined using the Arrhenius equation. For FL-*Pa*BphP-D, the activation energies for *k*_1_ and *k*_2_ are 2.44 and 5.11 kJ/mol, respectively. These values lead to the unexpected conclusion that the photochemical state of the sister protomer in the dimer influences the activation energy of the reaction. In contrast, PSM-*Pa*BphP-M shows a significantly higher activation energy of 15.21 kJ/mol for *k*_1_, suggesting that even the quaternary structure of the protein impacts the activation energy.

Our kinetic analysis shows that the transition rate (*k*_1_) from PrPr to PrPfr is about 2-3 times faster than the transition rate (*k*_2_) from PrPfr to PfrPfr. This suggests that a significant portion of the PrPfr intermediate state is formed during the intermediate phase of dark reversion in FL-*Pa*BphP-D (Fig. 4(f)). More importantly, this shows that substantial allosteric interactions may occur between the two protomers in the dimer, as the dark reversion time differs depending on whether both monomers are in the Pr state or if one is in the Pfr state. This points to an as-yet undescribed interaction between the two protomers in the dimer.

## DISCUSSION

Our kinetic analysis of *Pa*BphP provides the first direct demonstration of a PrPfr heterodimer intermediate during the thermal reversion of bacterial phytochromes. By analyzing kinetic models in a series of experiments with varying initial concentrations of Pr and Pfr, we show that a two-step mechanism accurately describes the dark reversion pathway. Only this model yields rate constants that are independent of the initial Pr:Pfr ratio

This finding mirrors observations made for plant phytochromes, where dark reversion proceeds via a PrPfr heterodimer before returning to the Pr state. ^41,44^ *Pa*BphP being a bathy phytochrome exhibits a similar sequential interconversion pathway, but in the reverse direction, from Pr to Pfr.

To our knowledge and somewhat surprisingly, earlier studies on bacterial phytochromes have not considered the possibility of a PrPfr intermediate state ^22,28,37,52,57,61,35,45–47,51,58^. This may be due to the fact that, in canonical phytochromes, the initial state is typically a mixture of Pr and Pfr, making it difficult to assign a specific dark recovery pathway. In contrast, bathy phytochromes allow for the generation of nearly pure Pr states through far-red illumination, enabling the identification of a PrPfr intermediate state.

It is established that the Pfr form of *Pa*BphP represents the biochemical “off” state. ^37,52^ Huber *et al*. qualitatively correlated biochemical kinase activity, measured via radionucleotide assays, with the decaying Pr/Pfr ratio during dark reversion. ^52^ However, their study did not consider the potential presence of a mixed PrPfr heterodimer. This raises the question of whether the PrPfr state is also biochemically active and whether it might even substitute for PrPr as the functional state. Notably, in plant phytochromes, the hybrid PrPfr state is known to play a distinct functional role compared to the pure PrPr or PfrPfr states. ^41,56^ A definitive assignment of the PrPfr states function would require a quantitative analysis of the radionucleotide assays, a challenging task, as acknowledged by Huber *et al*.. Nonetheless, their assays show high biochemical activity after prolonged far-red illumination, ^52^, a condition in which the PrPr population is significantly higher than that of PrPfr (see Fig. 4(c) at *φ* = 1). We believe that PrPr is indeed kinase-active.

However, this line of reasoning does not rule out the possibility that the PrPfr state also exhibits biochemical activity.

Our temperature-dependent measurements reveal activation energies that are significantly lower than those typically reported for dark reversion in plant phytochromes. In studies on the effect of temperature on the dark reversion properties of *Arabidopsis thaliana* phyA, activation enthalpies of 92.9 and 74.5 kJ*·*mol^*−*1^ were reported for PfrPfr homodimers and PrPfr heterodimers, respectively. ^42^ Another study found that all four isoforms showed strong temperature dependence, with activation energies of 104, 62, 163, and 75.3 kJ*·*mol^*−*1^ for PhyA, PhyB, PhyC, and PhyE, respectively. ^43^ Furthermore, phosphorylation has been shown to modulate the dark reversion rate. ^55^

The unusually low activation barriers observed for the *Pa*BphP protein are difficult to explain solely by a steric model of C15=C16 double bond isomerization. An alternative chemical mechanism may be involved. In fact, ketoenol tautomerization of the D-ring in its Z-isomerized (Pr) state has been proposed to precede isomerization of the C15C16 bond. ^51^ The enol form lowers the bond order of the C15C16 bond, which would significantly reduce the energy of the transition state and could account for the unusually low activation energy observed in our measurements.

Interestingly, we observe distinct rates and activation energies for the two steps of dark reversion (Fig. 5), suggesting that the state of the sister monomer within the dimer influences the chromophores reversion reaction. This strongly indicates an allosteric interaction. Supporting this, the monomeric variant, which lacks a dimer interface, exhibits an even higher activation energy for dark reversion (Fig. 5b). A similar observation has been made for plant phytochromes, suggesting that the two-step mechanism is tightly regulated by the protein and likely optimized through evolution ^42^

**Figure 5:**
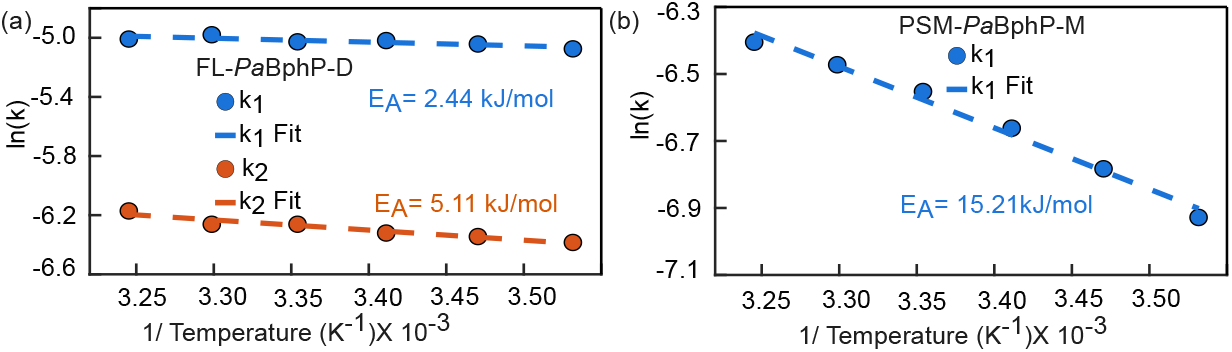
Arrhenius analysis shows notably low activation energies for the dark reversion of *Pa*BphP. Logarithmic variation of (a) fast (*k*_1_) and slow (*k*_2_) dark reversion rate constants with the inverse of temperature for FL-*Pa*BphP-D and (b) logarithmic variation of fast (*k*_1_) dark reversion rate constants with the inverse of temperature for PSM-*Pa*BphP-M. Activation energy is calculated using Arrhenius equation 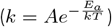

So far, allostery in bacterial phytochromes has primarily been discussed in the context of linking chromophore photoactivation to the regulation of biochemical output. However, the allosteric interaction observed here happens between the sister protomers within the dimer, suggesting the existence of a previously unidentified signaling pathway between the two protein subunits. Recent NMR investigations of the *D. radiodurans* phytochrome support this idea, revealing chemical shift changes that propagate from the chromophore to the dimer interface within the PAS/GAF domains ^62^. Further structural studies are needed to uncover the molecular basis of this signaling route.

## CONCLUSION

In this study, we have successfully demonstrated the transient accumulation of a mixed PrPfr hybrid state in the bacterial phytochrome *Pa*BphP during the dark reversion process. Our kinetic analysis indicates that both subunits within the dimeric phytochrome can exist in different photoisomeric states simultaneously during dark reversion, and that the hybrid state is transiently populated at high concentration. We also observe that a monomeric variant of *Pa*BphP follows a direct Pr-to-Pfr reaction scheme, which further supports the assignment of the PrPfr intermediate as a feature arising from the dimeric nature of the wild-type protein. Furthermore, the temperature-dependent kinetics reveal low activation energies for the dark reversion processes in both the dimeric and monomeric PaBphP, consistent with a previously proposed keto-enol tautomerization as a crucial step preceding dark reversion. The distinct kinetic behavior observed in the dimeric *Pa*BphP, specifically, the transient accumulation of the PrPfr hybrid state and the difference in activation energy, suggests the presence of allosteric regulation between the two subunits. We propose that this plays an important role in controlling the signaling output of bacterial phytochromes.

## ACKNOWLEDGEMENTS

The author acknowledge Göran Gustaffson foundations and Carl-Trygger Foundation for supporting this work. MM acknowledges the Swedish Research Council (Grant No. VR 2020-05403), the Swedish Society for Medical Research (S20-0156), and Harald and Greta Jeanssons Foundation Grant (J2021-0114).

## REFERENCES

[1] P. Hegemann, annurev.arplant, 2008, 59, 167–189.

[2] A. Möglich, X. Yang, R. A. Ayers and K. Moffat, annurev.arplant, 2010, 61, 21–47.

[3] N. C. Rockwell, Y.-S. Su and J. C. Lagarias, annurev.arplant, 2006, 57, 837–858.

[4] H. Wang and H. Wang, Mol. Plant., 2015, 8, 540–551.

[5] T. Lamparter, N. KrauSS and P. Scheerer, Photochem. Photobiol., 2017, 93, 642–655.

[6] J. J. Casal and R. A. Sánchez, Seed Sci. Res, 1998, 8, 317329.

[7] E. Monte, J. M. Tepperman, B. Al-Sady, K. A. Kaczorowski, J. M. Alonso, J. R. Ecker, X. Li, Y. Zhang and P. H. Quail, Proc. Natl. Acad. Sci. U.S.A, 2004, 101, 16091–16098.

[8] M. Endo, Y. Tanigawa, T. Murakami, T. Araki and A. Nagatani, Proc. Natl. Acad. Sci. U.S.A, 2013, 110, 18017–18022.

[9] J. A. Kyndt, T. E. Meyer and M. A. Cusanovich, Photochem. Photobiol. Sci, 2004, 3, 519–530.

[10] L. Wu, R. S. McGrane and G. A. Beattie, MBio, 2013, 4, 10–1128.

[11] R. K. Verma, A. Biswas, A. Kakkar, S. K. Lomada, B. B. Pradhan and S. Chatterjee, Cell Rep, 2020, 32, year.

[12] K. G. Chernov, T. A. Redchuk, E. S. Omelina and V. V. Verkhusha, Chem Rev, 2017, 117, 6423–6446.

[13] A. A. Kaberniuk, A. A. Shemetov and V. V. Verkhusha, Nat. Methods, 2016, 13, 591–597.

[14] A. T. Ulijasz and R. D. Vierstra, Curr. Opin. Plant Biol., 2011, 14, 498–506.

[15] A. Möglich, R. A. Ayers and K. Moffat, Structure, 2009, 17, 1282–1294.

[16] L. Aravind and C. P. Ponting, Trends Biochem. Sci., 1997, 22, 458–459.

[17] L.-O. Essen, J. Mailliet and J. Hughes, Proc. Natl. Acad. Sci. U.S.A, 2008, 105, 14709–14714.

[18] E. S. Burgie and R. D. Vierstra, Plant Cell, 2014, 26, 4568–4583.

[19] M. Legris, C. Klose, E. S. Burgie, C. C. R. Rojas, M. Neme, A. Hiltbrunner, P. A. Wigge, E. Schäfer, R. D. Vierstra and J. J. Casal, Science, 2016, 354, 897–900.

[20] G. Rottwinkel, I. Oberpichler and T. Lamparter, J. Bacteriol. Res, 2010, 192, 5124–5133.

[21] F. Velázquez Escobar, D. Buhrke, N. Michael, L. Sauthof, S. Wilkening, N. N. Tavraz, J. Salewski, N. Frankenberg-Dinkel, M. A. Mroginski, P. Scheerer et al., Photochem. Photobiol, 2017, 93, 724– 732.

[22] M. F. López, M. Dahl, F. V. Escobar, H. R. Bonomi, A. Kraskov, N. Michael, M. A. Mroginski, P. Scheerer and P. Hildebrandt, Phys. Chem. Chem. Phys., 2022, 24, 11967–11978.

[23] F. Velázquez Escobar, D. Buhrke, N. Michael, L. Sauthof, S. Wilkening, N. N. Tavraz, J. Salewski, N. Frankenberg-Dinkel, M. A. Mroginski, P. Scheerer et al., Photochem. Photobiol, 2017, 93, 724– 732.

[24] S. Westenhoff, P. Meszaros and M. Schmidt, Curr. Opin. Struct. Biol., 2022, 77, 102481.

[25] A. Björling, O. Berntsson, H. Lehtivuori, H. Takala, A. J. Hughes, M. Panman, M. Hoernke, S. Niebling, Henry, R. Henning, I. Kosheleva, V. Chukharev, N. V. Tkachenko, A. Menzel, G. Newby, D. Khakhulin, Wulff, J. A. Ihalainen and S. Westenhoff, Sci. Adv., 2016, 2, e1600920.

[26] M. Chenchiliyan, J. Kübel, S. A. Ooi, G. Salvadori, B. Mennucci, S. Westenhoff and M. Maj, J. Chem. Phys., 2023, 158, 085103.

[27] S. G. Sokolovski, E. A. Zherebtsov, R. K. Kar, D. Golonka, R. Stabel, N. B. Chichkov, A. Gorodetsky, I. Schapiro, A. Möglich and E. U. Rafailov, Biophys. J., 2021, 120, 964–974.

[28] Y. Yang, T. Stensitzki, L. Sauthof, A. Schmidt, P. Piwowarski, F. Velazquez Escobar, N. Michael, A. D. Nguyen, M. Szczepek, F. N. Brünig, R. R. Netz, M. A. Mroginski, S. Adam, F. Bartl, I. Schapiro, P. Hildebrandt, P. Scheerer and K. Heyne, Nat. Chem., 2022, 14, 823–830.

[29] H. Takala, P. Edlund, J. A. Ihalainen and S. Westenhoff, Photochem. Photobiol. Sci., 2020, 19, 1488– 1510.

[30] X. Yang, Z. Ren, J. Kuk and K. Moffat, Nature, 2011, 479, 428–432.

[31] X. Yang, J. Kuk and K. Moffat, Proc. Natl. Acad. Sci. U.S.A, 2009, 106, 15639–15644.

[32] J. R. Wagner, J. S. Brunzelle, K. T. Forest and R. D. Vierstra, Nature, 2005, 438, 325–331.

[33] L.-O. Essen, J. Mailliet and J. Hughes, Proceedings of the National Academy of Sciences, 2008, 105, 14709–14714.

[34] X. Yang, J. Kuk and K. Moffat, PNAS, 2008, 105, 14715–14720.

[35] W. Y. Wahlgren, E. Claesson, I. Tuure, S. Trillo-Muyo, S. Bódizs, J. A. Ihalainen, H. Takala and S. Westenhoff, Nat. Commun, 2022, 13, 7673.

[36] H. Li, E. S. Burgie, Z. T. K. Gannam, H. Li and R. D. Vierstra, Nature, 2022, 604, 127–133.

[37] S. Bódizs, P. Mészáros, L. Grunewald, H. Takala and S. Westenhoff, Structure, 2024, 32, 1952– 1962.e3.

[38] D. Buhrke, G. Gourinchas, M. Müller, N. Michael, P. Hildebrandt and A. Winkler, J. Biol. Chem., 2020, 295, 539–551.

[39] T. N. Malla, C. Hernandez, S. Muniyappan, D. Menendez, D. Bizhga, J. H. Mendez, P. Schwander, E. A. Stojkovi and M. Schmidt, Sci. Adv, 2024, 10, eadq0653.

[40] E. Schäfer and W. Schmidt, Planta, 1974, 116, 257–266.

[41] J. Brockmann, S. Rieble, N. Kazarinova-Fukshansky, M. Seyfried and E. Schäfer, Plant Cell Environ., 1987, 10, 105–111.

[42] L. Hennig and E. Schäfer, J. Biol. Chem., 2001, 276, 7913–7918.

[43] E. S. Burgie, Z. T. K. Gannam, K. E. McLoughlin, C. D. Sherman, A. S. Holehouse, R. J. Stankey and R. D. Vierstra, Proc. Natl. Acad. Sci. U.S.A, 2021, 118, e2105649118.

[44] W. Schmidt and E. Schäfer, Planta, 1974, 116, 267–272.

[45] C. Huber, M. Strack, I. SchultheiSS, J. Pielage, X. Mechler, J. Hornbogen, R. Diller and N. Frankenberg-Dinkel, J. Biol. Chem., 2024, 300, 107148.

[46] A. Kraskov, A. D. Nguyen, J. Goerling, D. Buhrke, F. Velazquez Escobar, M. Fernandez Lopez, Michael, L. Sauthof, A. Schmidt, P. Piwowarski, Y. Yang, T. Stensitzki, S. Adam, F. Bartl, I. Schapiro, K. Heyne, F. Siebert, P. Scheerer, M. A. Mroginski and P. Hildebrandt, Biochemistry, 2020, 59, 1023– 1037.

[47] G. Merga, M. F. Lopez, P. Fischer, P. Piwowarski,. Nogacz, A. Kraskov, D. Buhrke, F. V. Escobar, N. Michael, F. Siebert, P. Scheerer, F. Bartl and P. Hildebrandt, Phys. Chem. Chem. Phys., 2021, 23, 18197–18205.

[48] H. Takala, H. Lehtivuori, H. Hammarén, V.P. Hytönen and J. A. Ihalainen, Biochemistry, 2014, 53, 7076–7085.

[49] E. S. Burgie, Z. T. K. Gannam, K. E. McLoughlin, C. D. Sherman, A. S. Holehouse, R. J. Stankey and R. D. Vierstra, PNAS, 2021, 118, e2105649118.

[50] J.-H. Jung, M. Domijan, C. Klose, S. Biswas, D. Ezer, M. Gao, A. K. Khattak, M. S. Box, V. Charoensawan, S. Cortijo, M. Kumar, A. Grant, J. C. W. Locke, E. Schäfer, K. E. Jaeger and P. A. Wigge, Science, 2016, 354, 886–889.

[51] F. Velazquez Escobar, P. Piwowarski, J. Salewski, N. Michael, M. Fernandez Lopez, A. Rupp, B. M. Qureshi, P. Scheerer, F. Bartl, N. Frankenberg-Dinkel, F. Siebert, M. Andrea Mroginski and P. Hildebrandt, Nat. Chem., 2015, 7, 423–430.

[52] C. Huber, M. Strack, I. SchultheiSS, J. Pielage, X. Mechler, J. Hornbogen, R. Diller and N. Frankenberg-Dinkel, J. Biol. Chem., 2024, 300, 107148.

[53] H. Takala, A. Björling, M. Linna, S. Westenhoff and J. A. Ihalainen, J. Biol. Chem., 2015, 290, 16383– 16392.

[54] W. Schmidt and E. Schäfer, Planta, 1974, 116, 267–272.

[55] M. Medzihradszky, J. Bindics,. Ádám, A. Viczián, Klement, S. Lorrain, P. Gyula, Z. Mérai, C. Fankhauser, K. F. Medzihradszky, T. Kunkel, E. SchÄfer and F. Nagy, The Plant Cell, 2013, 25, 535–544.

[56] C. Klose, F. Venezia, A. Hussong, S. Kircher, E. Schäfer and C. Fleck, Nat. Plants, 2015, 1, 15090.

[57] H. Takala, A. Björling, M. Linna, S. Westenhoff and J. A. Ihalainen, J. Biol. Chem., 2015, 290, 16383– 16392.

[58] K. D. Piatkevich, F. V. Subach and V. V. Verkhusha, Nat. Commun., 2013, 4, 2153.

[59] A. Kraskov, A. D. Nguyen, J. Goerling, D. Buhrke, F. Velazquez Escobar, M. Fernandez Lopez, N. Michael, L. Sauthof, A. Schmidt, P. Piwowarski, Y. Yang, T. Stensitzki, S. Adam, F. Bartl, I. Schapiro, K. Heyne, F. Siebert, P. Scheerer, M. A. Mroginski and P. Hildebrandt, Biochemistry, 2020, 59, 1023– 1037.

[60] M. E. Auldridge, K. A. Satyshur, D. M. Anstrom and K. T. Forest, J. Biol. Chem., 2012, 287, 7000–7009.

[61] R. Tasler, T. Moises and N. Frankenberg-Dinkel, The FEBS Journal, 2005, 272, 1927–1936.

[62] L. Isaksson, E. Gustavsson, C. Persson, U. Brath, L. Vrhovac, G. Karlsson, V. Orekhov and S. Westenhoff, Structure, 2020, 29, 151–160.e3.

